# Boolean Implication Analysis Improves Prediction Accuracy of In Silico Gene Reporting of Retinal Cell Types

**DOI:** 10.1101/2020.09.28.317313

**Authors:** Rohan Subramanian, Debashis Sahoo

**Affiliations:** The International School Bangalore, Bengaluru, Karnataka, India; Department of Pediatrics, University of California San Diego, La Jolla, CA, USA; Department of Computer Science and Engineering, Jacobs School of Engineering, University of California San Diego, La Jolla, CA, USA

**Author notes:** **Corresponding Author**: **Debashis Sahoo, Ph.D.;** Assistant Professor, Department of Pediatrics, University of California San Diego; 9500 Gilman Drive, MC 0730, Leichtag Building 132; La Jolla, CA 92093-0831. : : **Email:**.

**Keywords:** Retina, Single-cell RNA sequencing, Pluripotent stem cells, Boolean analysis, Bioinformatics

## Abstract

The retina is a complex tissue containing multiple cell types that is essential for vision.
Understanding the gene expression patterns of various retinal cell types has potential applications in regenerative medicine. Retinal organoids (optic vesicles) derived from pluripotent stem cells have begun to yield insights into the transcriptomics of developing retinal cell types in humans through single cell RNA-sequencing studies. Previous methods of gene reporting have relied upon techniques in vivo using microarray data, or correlational and dimension reduction methods for analyzing single cell RNA-sequencing data in silico. Here, we present a bioinformatic approach using Boolean implication to discover retinal cell type-specific genes. We apply this approach to previously published retina and retinal organoid datasets and improve upon previously published correlational methods. Our method improves the prediction accuracy and reproducibility of marker genes of retinal cell types and discovers several new high confidence cone and rod-specific genes. Furthermore, our method is general and can impact all areas of gene expression analyses in cancer and other human diseases.

**Significance Statement:** Efforts to derive retinal cell types from pluripotent stem cells to the end of curing retinal disease require robust characterization of these cell types’ gene expression patterns. The Boolean method described in this study improves prediction accuracy of earlier methods of gene reporting, and allows for the discovery and validation of retinal cell type-specific marker genes. The invariant nature of results from Boolean implication analysis can yield high-value molecular markers that can be used as biomarkers or drug targets.

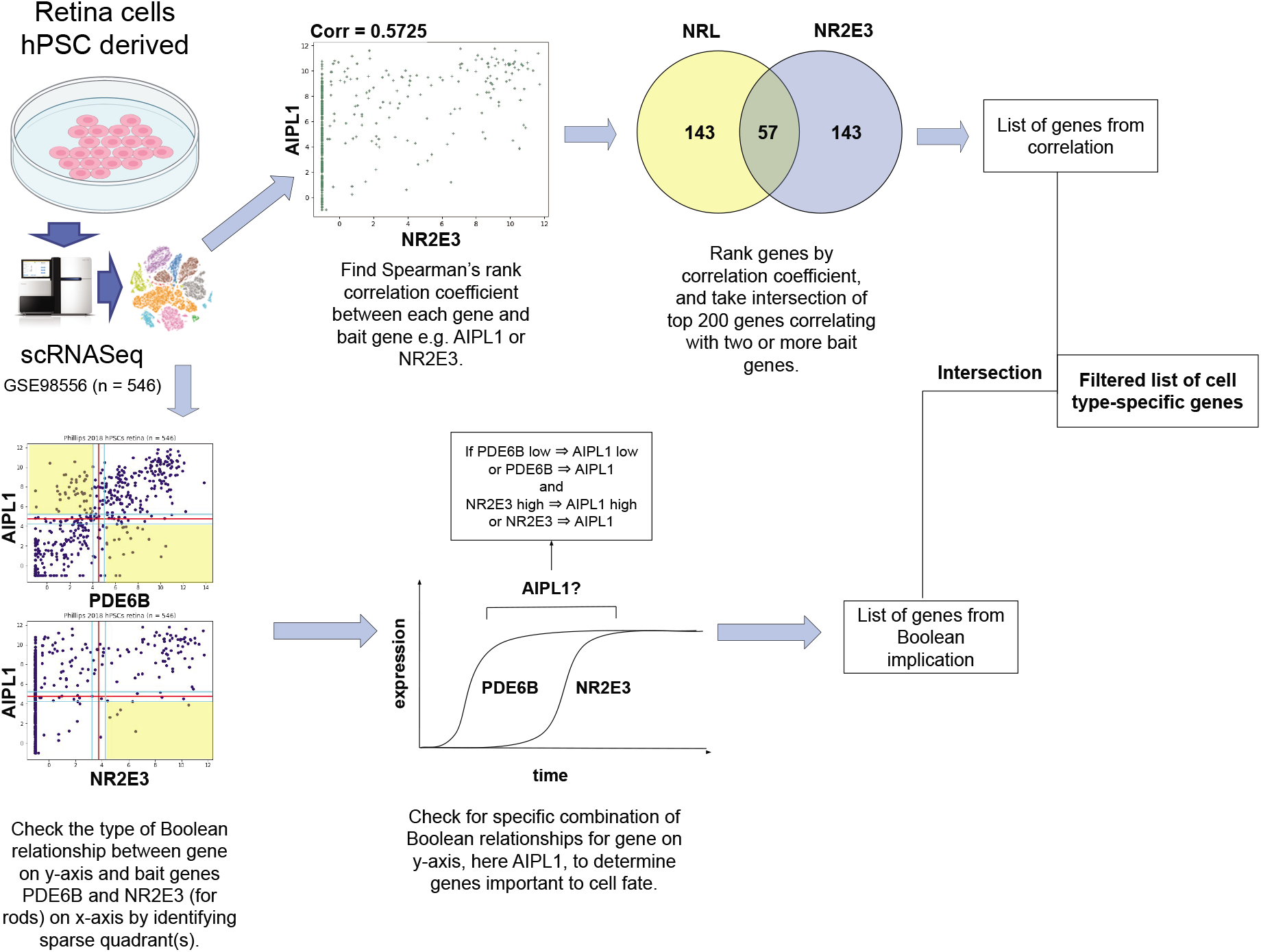

## Introduction

Characterization of retinal cell types is an important field of study with wide applications in ophthalmology and regenerative medicine. With the advent of single cell RNA-sequencing (scRNA-seq), methods for gene reporting in silico can yield valuable insights into genes that are important in determining cell fate.^1^ Human pluripotent stem cells (hPSCs) can be used to generate retinal cell types in vitro with potential applications to cure age-related macular degeneration, retinitis pigmentosa and other retina-related causes of blindness. However, gene reporting and characterization of these cell types is difficult as they differentiate asynchronously in complex cultures.^2^ Furthermore, there is a lack of human datasets. We propose using Boolean implication analysis to improve the prediction accuracy of existing correlational methods for in silico gene reporting.

### Previous Methods In Vivo and In Vitro

One of the most common methods to study the effect of key genes on retinal development is the use of genetically modified “knockout” murine models, which are frequently used to validate differentially expressed genes from microarray data.^3–20^ Fluorescent gene reporter lines are widely used to check for gene expression in single cells, or purified populations of a single cell type.^2, 21–25^ Bulk RNA sequencing (RNA-seq) has helped define the transcriptomes of larger populations of retinal cell types.^3, 9, 14, 17, 21, 24, 26–35^ To study the characteristics of isolated cells or droplets, flow cytometry was formerly a major method.^36, 37^ Single-cell RNA sequencing (scRNA-seq) is increasingly common today and is one the most detailed methods to profile transcriptomes of retinal cell types and subtypes.^2, 8, 13, 22, 38–48^

Most studies on retinal cell types have relied upon murine models, but many increasingly study human donor retinas^6, 30, 31, 48–50^, especially in order to profile retinal disease.^31, 43, 50–53^ Glaucoma, age-related macular dystrophy and retinal light damage have also been studied in murine models.^7, 14, 29, 34, 35, 54, 55^ Some studies have grown cell lines in vitro from fetal retina^49, 56^, whereas other have used human pluripotent, induced pluripotent or embryonic stem cells to generate purified cell populations or retinal organoids.^2, 3, 8, 28, 38, 57–59^ In order to study the development of retinal cell types over time, the lineage of stem cell progeny^58^ and time course data from different time points (using PCR and RNA-seq) have been investigated.^39, 41, 54^

### Previous Methods In Silico

Differential expression analysis is the most common method to identify retinal cell type-specific genes and biomarkers from microarray, RNA-seq and scRNA-seq data.^10, 13, 14, 17, 24, 29–31, 39, 41, 46, 47, 53, 56, 59^ In single-cell analysis, dimension reduction through Principal Component Analysis to reduce the size of data and allow visualization is often performed before hierarchical clustering identify cell clusters.^2, 7, 30, 41, 42, 49, 56, 60^ Cell clusters can be assigned to different cell types or subtypes based on the expression of key marker genes.^48^ AI-guided identification of cell clusters has recently been investigated.^61^

scRNA-seq data provides opportunities for in depth analysis of the transcriptome of individual cells, and subsequent characterization of cell types, subtypes and regions of retina. However, scRNA-seq data is highly noisy, and contains large numbers of zeroes, among which true and false negatives are indistinguishable. Many of these zeroes are dropouts, caused by a failure to capture or amplify a transcript.^62^

Most studies up to date have been highly dependent on cell clustering, which is not always achievable, especially in datasets containing immature or developing cells.^1^ Pseudo-time analysis, which aims to sort cells by their developmental stage, has been applied to retinal organoids, and takes into account transitory states rather than discrete clusters.^38^ However, this approach is hindered by asynchronous differentiation of cell types in retina.^63^ Correlational methods for ranking gene expression are also widely used, bypassing the need to discover cell clusters and identifying co-expressed genes in complex cultures, including developing retinal organoids.^2,8,23,27,49,64^

Identifying relationships between genes has led towards broader goals of graph^47, 60^ and networkbased analysis.^9, 10, 17, 25, 27, 31, 60, 65^ Gene expression networks can be used to identify transitions between phenotypes and disease states, paving the way for clinical target identification. Correlational analysis is traditionally used to derive co-expression networks, and knockout murine models are used to directly investigate the effect of one gene’s absence of others. However, the symmetric nature of correlation can lead to loss of valuable information and does not provide insight into the expression of genes over time. Bayesian networks of gene regulation and expression in the retina mainly identify transcription factors and their targets.^60, 66^ Hence, the motivation of our work was to develop a universally applicable state of the art method that filtered out noise, could be applied to a wide variety of datasets and lent insight into gene expression over differentiation.

### A Boolean Approach

Boolean logic is a simple mathematical relationship between two values such as high/low or 1/0. We propose using Boolean implication (“if-then” relationships) to study the dependency between genes from scRNA-seq data. Research by Sahoo et al. has shown that analysis of Boolean implication relationships is better at filtering out noise than correlational approach.^67^ Analysis of Boolean implication lends insight into asymmetric relationships disregarded by correlation.

While Boolean implication, like correlation, does not imply causation, asymmetric Boolean relationships can be thought of in terms of subsets. For example, the relationship Gene A high ⇒ Gene B high indicates that all cells with Gene B high are a subset of those with Gene A high. This allows for analysis of developmentally regulated genes using Boolean implication, first pioneered in the MiDReG tool published by Sahoo et al 2010.^68^

In previous research, Boolean methods have led to the discovery of prognostic biomarkers for bladder and colon cancer.^69–71^ These methods have also led to characterization of hematopoietic stem cells and identification of B and T cell precursors.^72, 73^ Our methods have not previously been applied to stem cell-derived retinal cell types, but have yielded insights into changes in transcriptional profiles of healthy retina and retinoblastoma.^74^

The StepMiner and BooleanNet algorithms were developed for microarray data by Sahoo et al. 2008 to identify Boolean implication relationships between genes, but have since been applied to a wide variety of high-throughput data, such as RNA-seq, and scRNA-seq.^68, 75, 76^

## Methods

### Data Normalization and Annotation

We applied log2(v+1) transformation to TPM values from the 546 sequenced cells of the Phillips 2018 dataset (GSE98556, n = 546), as the log transformed RNA-seq data is closer to a normal distribution. Cells were annotated with clinical characteristics, and data were uploaded to Hegemon. In the Hegemon online tool, scatter plots between genes are generated, with each point representing the expression level of the genes in a single cell.^67–71^

### Discovering Boolean Implications

#### StepMiner Algorithm

The StepMiner algorithm identifies thresholds to convert continuous expression values into discrete values by fitting a step function to sorted values. A step can be defined as the sharpest increase in sorted gene expression values over an interval. Having identified a threshold *t*, gene expression values greater than *t + 0.5* are considered high, and those below *t - 0.5* are considered low. Those between *t + 0.5* and *t - 0.5* are considered intermediate. These thresholds are used to divide the plot into four quadrants.^67, 77^

#### BooleanNet Algorithm

The BooleanNet algorithm identifies the type of Boolean implication relationship by identifying the sparse quadrant(s) using a statistic *S* and likelihood error rate *p*. There are six types of Boolean implication relationship: high ⇒ high, low ⇒ low, high ⇒ low, low ⇒ high, equivalent and opposite. The first four are asymmetric and have only one sparse quadrant. The latter two are symmetric and have two sparse quadrants. Further information can be found in **Fig. S1**.^67, 77^

#### Thresholds for Analysis

Thresholds for S and p are applied to adjust the sensitivity of the analysis. While S > 3 and p < 0.1 are generally considered for microarray data, we decreased the threshold for S to 2.5 and increased the threshold for p to 0.25 for rods and 0.35 for all other analysis to account for the larger amount of noise in scRNA-seq data.

### Boolean Approach to In Silico Gene Reporting of Retinal Cell Types

We propose using the method described in **Fig. 1** for in silico gene reporting of retinal cell types. We require two or more known genes for each cell type called “bait genes”. We searched for genes which had a low ⇒ low or equivalent Boolean relationship with the first bait gene and high ⇒ high or equivalent Boolean relationship with the second bait gene.

**Figure 1.**
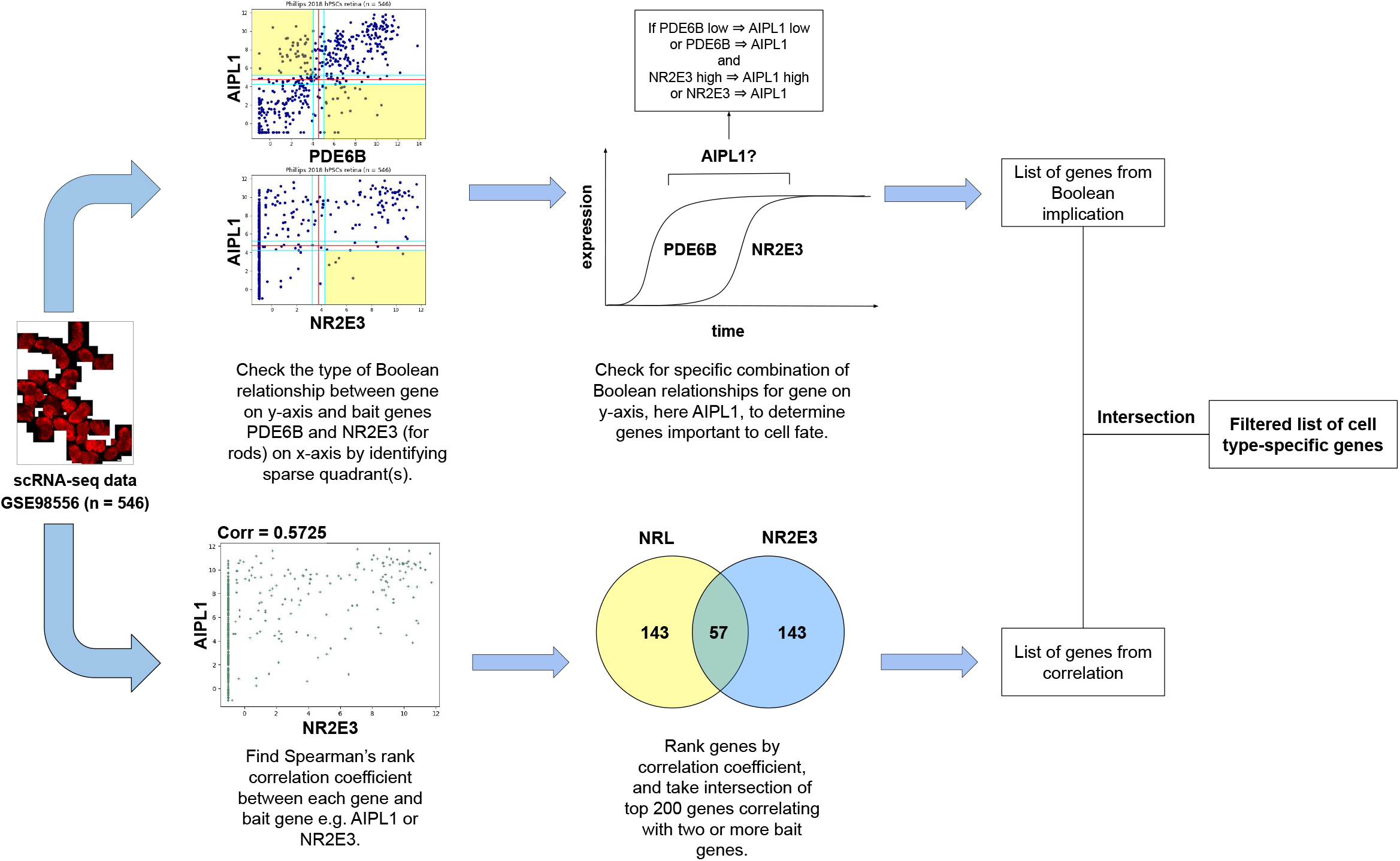
Schematic Algorithm. Schematic algorithm to discover cell type-specific genes from scRNA-seq data by combining correlational and Boolean implication analysis. Boolean implication analysis uses one general and one specific bait gene to identify cell type-specific biomarkers. Spearman’s rank correlation coefficient analysis (SRCCA) uses one or more genes specific to a cell type as bait genes to identify other genes expressed in the same cell type. Boolean analysis is directly compared to SRCCA and improvement is tested using two proportion Z-test.

This specific combination of Boolean relationships is akin to searching for genes which have an impact on cell fate. If a gene passes this analysis, the set of cells where Gene X is low is a subset of the cells where the first bait gene is low, and the set of cells where Gene X is high is a subset of the cells where the second bait gene is high. This method can allow us to infer genes which are expressed after the first bait gene, and before the second bait gene. Hence, the choice of bait genes plays an important role in determining the results. We chose bait genes which led to shorter gene lists compared to SRCCA, with a greater number of known markers of five retinal cell types. These were selected and verified from previous literature on rod and cone photoreceptors^6, 78^, retinal progenitor cells (RPCs)^79, 80^, retinal ganglion cells (RGCs)^23, 24^ and retinal pigment epithelium (RPE)^81–84^.

More than two bait genes can be considered by searching for high ⇒ high, low ⇒ low or equivalent Boolean relationships in two out of three bait genes instead of one out of two. This allows for combination of multiple cell type-specific marker genes in the analysis.

### Spearman’s Rank Correlation Coefficient

Spearman’s rank correlation coefficient (SRCC) is a nonparametric measure of the association between two ranked variables. We reviewed and reproduced the approach of Phillips et al. 2018, called Spearman’s rank correlation coefficient analysis (SRCCA). The correlation coefficient between bait genes and all other genes are found and ranked. Then, the intersection between the top 200 correlating genes with each bait gene is taken.^2^

We combined both methods by taking the interaction of gene lists derived from both methods, hence filtering the list of correlating genes using Boolean implication as shown in **Fig. 1**. All analysis was performed using the Hegemon website, in Python 3 using the HegemonUtil tools and in R version 4.0.1.

### Quantification of Results

Results were independently validated through differential expression. We evaluated whether genes were differentially expressed between rods and cones, and between photoreceptors and non-photoreceptor retinal cell types.

We selected and processed several validation datasets. Two were bulk RNA-seq datasets containing purified retinal cell types from Mus musculus: Hartl 2017 (GSE84589, n = 14) and Sarin 2018 (GSE98838, n = 22).^40, 46^ The third was a similar human retina scRNA-seq dataset, Voigt 2020 (GSE130636 and GSE142449, n = 20,797).^48^

Using validation datasets with purified cell types, we checked for differential expression between retinal cell types by performing a one-tailed Welch’s t-test between the groups of cells to determine whether there was a statistically significant difference between the means of the two groups. Using this method, we could evaluate the proportion of genes which were specific to the cell type in question, expressed equally throughout the retina, and expressed in a different, nontarget cell type.

To evaluate the reproducibility of the genes, we directly repeated the analysis in GSE130636 and GSE142449 using common bait genes for SRCCA and Boolean implication. We compared the proportion of genes discovered by using a two proportion Z-test.

## Results

### Boolean Implication Enables Identification of Cell Type Specific Genes like SRCCA

Boolean Implication analysis explores both symmetric and asymmetric relationship between genes whereas SRCCA only focuses on symmetric relationships. We hypothesize that application of asymmetric Boolean implication relationships may improve the accuracy of cell types specific genes identification (**Fig 1**).

Application of Boolean implication analysis led to shorter lists of genes compared to SRCCA **(Fig. 2)**. Selecting bait genes is crucial for both SRCCA and Boolean analysis. For Boolean analysis, a general marker and a more specific marker are ideal candidates. However, SRCCA relies only on specific bait genes. Because of these differences in specificity, we chose different set of bait genes for Boolean analysis from known marker genes for each retinal cell type.

**Figure 2.**
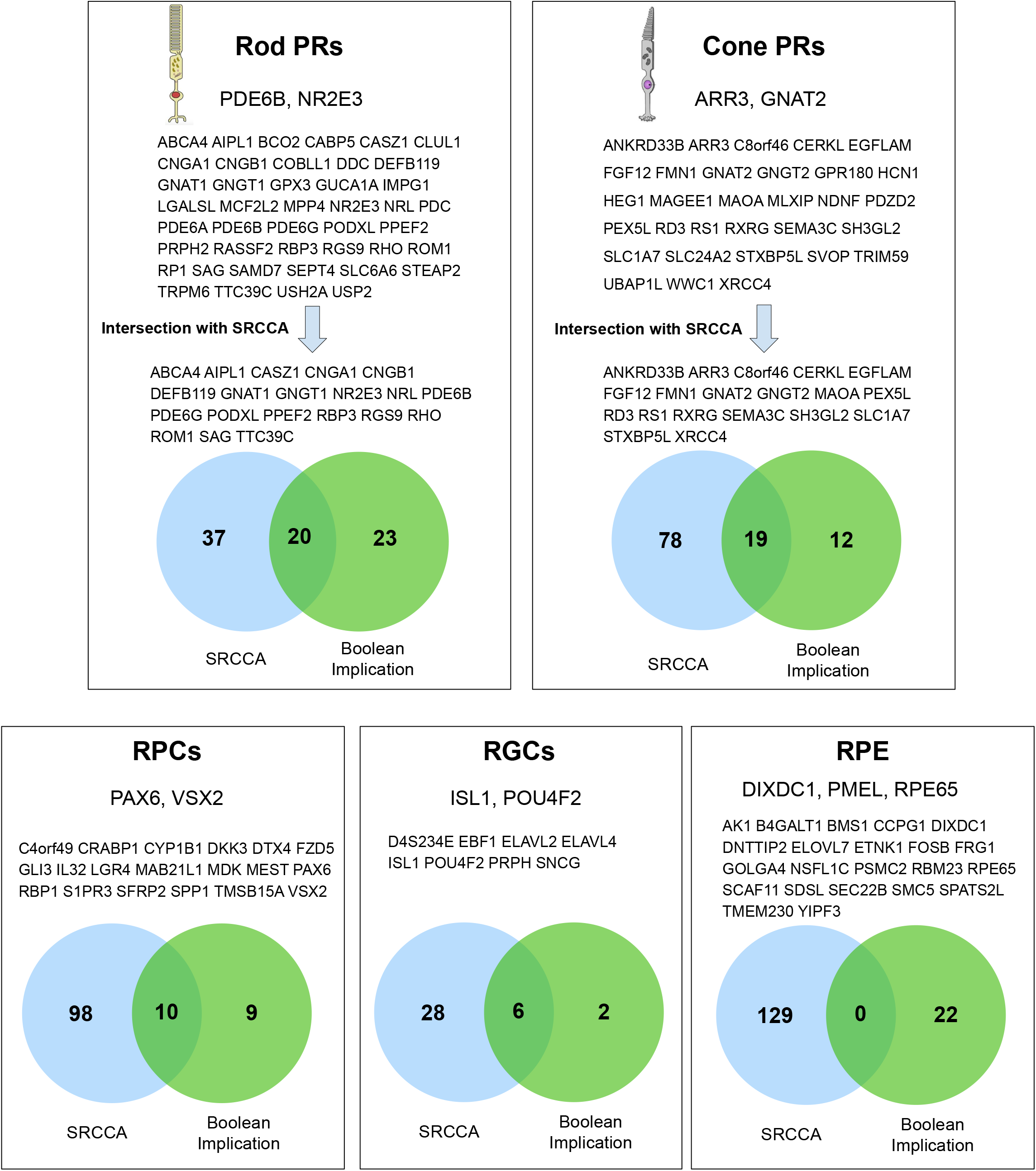
Results. Results of Boolean implication analysis of Phillips 2018 scRNA-seq dataset using two or more bait genes, for 5 retinal cell types. **Abbreviations**: SRCCA - Spearman’s Rank Correlation Coefficient Analysis: PR - photoreceptors; RPC - retinal progenitor cell; RPE - retinal pigment epithelium; RGC - retinal ganglion cell.

Application of Boolean analysis for gene reporting of photoreceptors led to longer lists of genes than other cell types. The largest intersection between SRCCA and Boolean implication was observed in rod photoreceptors. The number of genes from Boolean implication in other retinal cell types such as RGCs, RPCs and RPE was far lower than photoreceptors.

For RPE, three bait genes were chosen due to the excessively small number of genes obtained from two bait genes. This is likely to be due to the smaller number of cells from these types present in the retina, compared to photoreceptors. The complete absence of intersection between genes from SRCCA and Boolean in RPE could also be explained by the very small number of RPE cells present in optic vesicle cultures produced by the method used by Phillips et al. 2018.

### Filtering SRCCA using Boolean Implication Improves Prediction Accuracy

We independently validated the genes from SRCCA and Boolean implication using bulk RNA-seq datasets with purified retinal cell types. In **Fig. 3B**, there is a visible improvement in proportion of rod-specific genes while taking the intersection of SRCCA and Boolean implication. Similarly, the majority of SRCCA genes not present in Boolean implication were not specific to rods, or specific to cones. We were able to show a statistically significant improvement in the proportion of rod PR-specific genes by filtering correlating genes using Boolean implication. The proportion of genes rod-specific genes from SRCCA, 29 out of 56 (0.517), was improved to 16 out of 19 (0.842) by filtering using Boolean implication. This proportion was shown to be statistically significant by performing a two-proportion Z-test, returning a p-value of 0.013.

**Figure 3.**
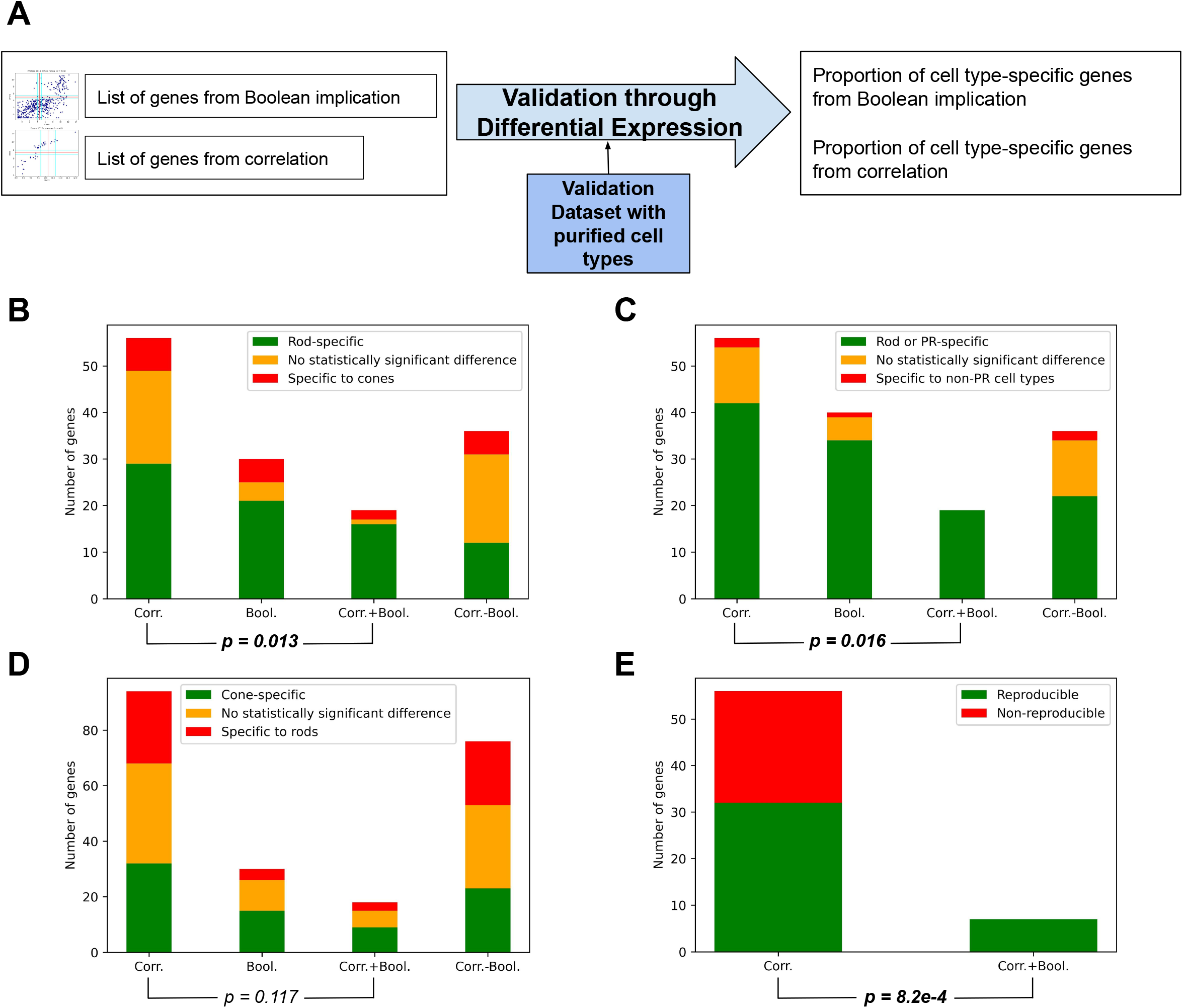
Independent Validation of Results. **(A):** Validation bulk RNA-seq datasets such as GSE84589 containing purified rods and cones from Mus musculus were used to validate rod and cone gene lists through differential expression. **(B):** Rod cell type-specificity of rod gene lists from 4 methods: Boolean implication, SRCCA, SRCCA filtered using Boolean implication and SRCCA without Boolean implication. **(C):** Photoreceptor-specificity of rod gene lists from 4 methods. **(D):** Cone cell type-specificity of rod gene lists from 4 methods. **(E):** Proportion of genes from SRCCA and SRCCA filtered using Boolean implication using a common set of bait genes CRX, GNB3 and GNAT2 directly reproducible in GSE130636 and GSE142449. **Abbreviations:** Corr. - Correlation; Bool. - Boolean; SRCCA - Spearman’s rank correlation coefficient analysis. **Note:** P-values are from two-proportion Z-test between proportion of cell type-specific genes in lists from SRCCA and SRCCA filtered using Boolean implication.

**Figure 4.**
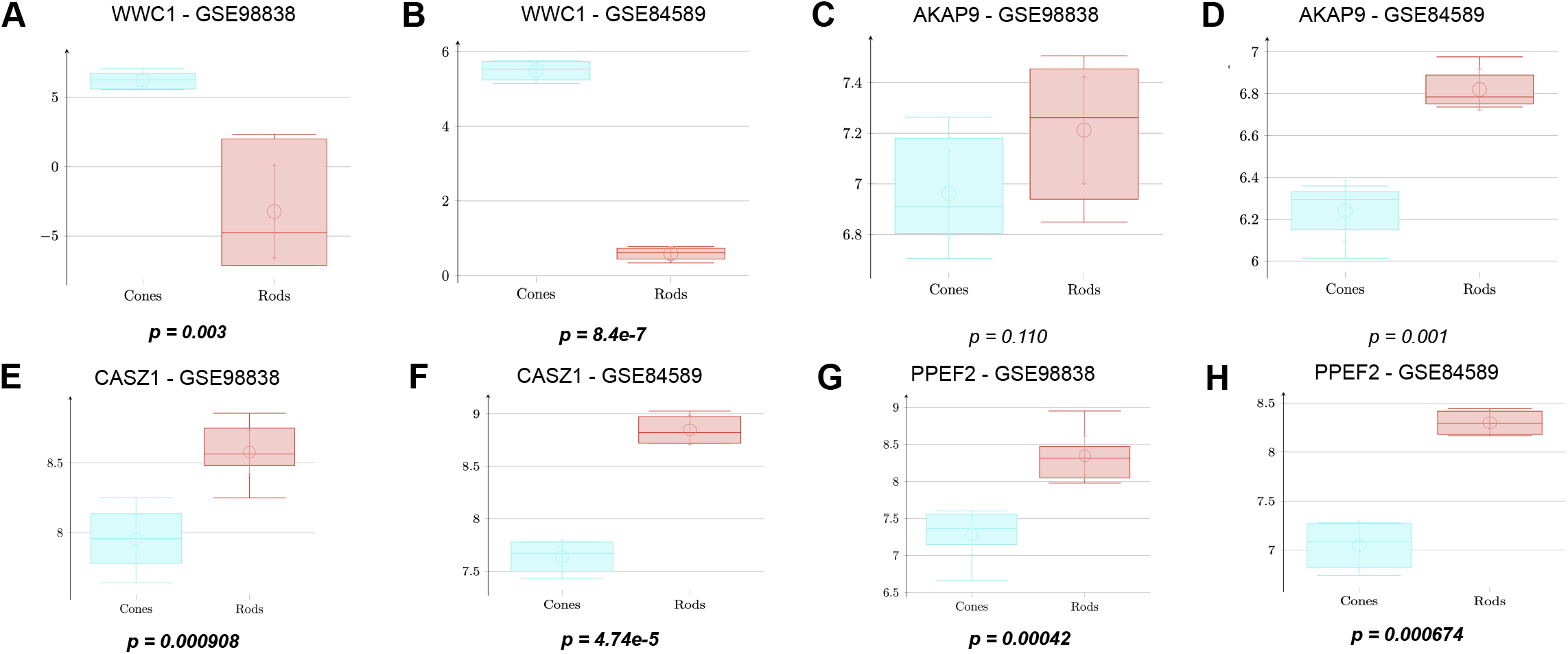
Specific Examples. Analysis of high confidence candidate PR genes from Boolean implication analysis and SRCCA. **(A-B):** High confidence cone PR gene WWC1 from Boolean implication analysis shows statistically significant overexpression in cones compared to rods in both datasets. **(C-D):** High confidence cone PR gene AKAP9 from SRCCA does not show cone specificity in either dataset. Cone PR group labelled in blue and rod PR group labelled in red on boxplots. **(E-H):** High confidence rod PR genes CASZ1 and PPEF2 from Boolean implication analysis show statistically significant overexpression in rods compared to cones in both datasets. **Note**: p-values reported from Welch’s t-test (unequal variances). **Abbreviations:** PR - photoreceptor; SRCCA - Spearman’s rank correlation coefficient analysis.

Similarly, as shown in **Fig. 3C**, we were able to show a statistically significant improvement in photoreceptor-specificity of the rod genes using the combined correlational and Boolean approach. All 19 genes obtained by filtering SRCCA using Boolean implication were photoreceptor-specific, and the p-value from the two-proportion Z-test was 0.016.

As seen in Fig. 3D, prediction accuracy of both SRCCA and Boolean analysis was lower in cone photoreceptors. The proportion of cone-specific genes, 15 out of 30 (0.500), was still highest in Boolean implication. Here, the prediction accuracy of Boolean methods alone was not improved by taking the intersection with SRCCA. However, this result could not be shown to be statistically significant due to the larger number of total genes in SRCCA. Hence, we sought an additional method of evaluation for cone photoreceptors with better scope for comparison.

### Filtering SRCCA using Boolean Implication Improves Reproducibility

We also performed SRCCA and Boolean implication analysis for cone photoreceptors using the same three bait genes: CRX, GNB3 and GNAT2. (**Fig. S2**) One major limitation of existing literature on characterization of retinal cell types is the lack of reproducibility of purported marker genes across datasets. Hence, we directly repeated the analysis in GSE130636 and GSE142449, which is also scRNA-seq of human retina. The main difference between this dataset and the Phillips dataset is the larger size (20,797 vs. 546 cells), and the larger proportion of adult, tissue-derived cells.

**Fig. 3E** displays the results from this method of quantification. Combining correlational and Boolean implication using common bait genes yielded highly reproducible results, as all 7 genes were reproduced. The two-proportion Z-test also returned a statistically significant p-value of 0.00082.

### Boolean Implication Improves Prediction Accuracy of Novel High Confidence Genes

Considering the overall improvement in prediction accuracy through Boolean implication analysis, we also investigated several specific examples of new discoveries through this method.

Novel high confidence genes are an important contribution of gene reporting methods in silico. Identification of high confidence markers of retinal cell types using SRCCA alone may be arbitrary, but we show that Boolean implication can lend greater insight into the cell typespecific genes.

Boolean implication analysis identified WWC1 (WW domain containing protein-1) as a novel high confidence cone photoreceptor gene. This was validated independently in GSE84589 and GSE98838, with statistically significant overexpression in cone photoreceptors. (**Fig. 3A-B**) WWC1 has been described to have a broad function in the brain and memory by previous studies.^85, 86^

Boolean implication analysis of rods also identified two novel rod-specific genes: CASZ1 (Castor zinc finger 1) (**Fig. 3E-F**) and PPEF2 (Protein Phosphatase with EF-Hand Domain 2) (**Fig. 3G-H**). These showed rod specificity in both validation datasets. CASZ1 is known to play a role in cell differentiation, and may hence play a significant role in influencing rod cell fate. ^87^ PPEF2 has been documented in rods before, but has had several conflicting studies on its importance in rods.^88, 89^ This is the first documentation of its rod-specific function in human or hPSC-derived retina. Boolean implication analysis has shed light on potential novel markers of cone and rod photoreceptors.

Boolean implication analysis refuted AKAP9 (A-kinase anchoring protein-9), identified to be a high confidence cone photoreceptor gene by Phillips et al. 2018 based on the results from SRCCA. **Fig. 3C-D** show that it is not differentially expressed in cones, and may be more rod-specific as per GSE84589.

## Discussion

Boolean methods improved upon correlational methods by filtering out noise and identifying asymmetric relationships that lend insight into the specificity of genes. Filtering correlating genes led to a statistically significant improvement in rod and photoreceptor-specificity for rod genes, and reproducibility for cone photoreceptor genes. Hence, we have shown that a combination of Boolean implication analysis and SRCCA improves the prediction accuracy of in silico gene reporting of retinal cell types.

Boolean implication analysis provided more accurate insight into high confidence genes, and led to the identification of WWC1 as a novel marker gene for cone photoreceptors. Previous attempts to identify high confidence genes from extensive gene lists obtained through SRCCA alone have no way to distinguish between noise and true cell type-specific genes. The asymmetric nature of Boolean relationships allows us to determine whether a gene is expressed more generally or specifically, which is not present in correlation.

Another advantage of Boolean implication is that the analysis can always be performed over the entire dataset. Boolean implication relationships between genes are best visible when there is a greater diversity of cell types, including those not expressing the gene. However, SRCCA generally requires the operator to choose a specific subset of the data (e.g. day 70) on which to perform the analysis, based on whether the cell type in question is present at that developmental stage. This choice has a significant effect on the result of SRCCA, and an inept choice of the subset may lead to false associations not generalizable over larger datasets. This issue can be solved using Boolean implication.

However, Boolean implication analysis was also not entirely free from error. The main source of error appears to be the dropouts, which lead to a greater density of points in quadrants a10, a00 and a01 in many cases. This, along with the slightly relaxed thresholds adapted for scRNA-seq, led to false discoveries of Boolean relationships. This issue likely reduced the improvement in quality of analysis in cone photoreceptors. Even so, a combination of correlational and Boolean implication analysis could lead to completely error-free results in some cases. (**Fig. 3C**)

The method of independent validation considered several datasets to evaluate specificity and reproducibility. These high-quality datasets provided reliable results for most genes, as human and mouse retina are very similar. However, it was not infallible due to small variations between the species. We compared the results in mouse datasets with the Kim 2019 dataset (GSE119343, n = 1346) containing cone-enriched optic vesicles. There, we found small differences in expression patterns in the mouse vs. human retina, such as CERKL, a gene specific to human cone photoreceptors, but expressed in both cone and rod photoreceptors in mice.

There were differences in the performance of our methods between different cell types. In cell types present in smaller numbers in the retina, we can observe that the number of genes from Boolean analysis alone and combined with SRCCA is also smaller. The analysis performed best in rods, the most numerous neural retina cell type.^90^ In RPE, which is rarely present in the optic vesicle culture protocol employed by Phillips et al. 2018, there was no intersection between Boolean analysis and SRCCA, indicating that the results in that case are likely to be mainly noise. However, there is no link between the number of genes obtained from SRCCA and the population of cell type, as a result of always considering a fixed number of top correlating genes. Hence, Boolean analysis can lend insight into the cell types for which the data is comprehensive enough to provide accurate resolution.

Boolean implication analysis provides all the advantages offered previously by SRCCA including efficiency, ability to combine multiple bait genes, and improved prediction accuracy compared to earlier methods. Our method can allow researchers to analyze single-cell data even when cell clusters cannot be identified, a common issue in datasets containing developing cells. Combining both methods provides statistically significant improvements in specificity and reproducibility of genes. Boolean implication can be easily inferred from scatter plots on the Hegemon online tool, making it an intuitive option for biologists and computer scientists alike.^76^

## Conclusion

In this work, we have developed a novel approach for analysis of scRNA-seq data based on Boolean implication. We have shown a statistically significant improvement in the prediction accuracy and reproducibility of retinal cell-type specific genes, as compared to earlier approaches based solely on correlation. Application of our method to retinal organoid datasets identified novel high confidence cell type-specific genes such as WWC1 for cones and CASZ1 and PPEF2 for rods. This Boolean approach allows for analysis and characterization of cell types in complex cultures, even when cell clustering cannot be achieved. Considering asymmetric relationships has allowed us to effectively filter out noise, lending insight into genes with potential importance in regenerative medicine.

## Author contributions

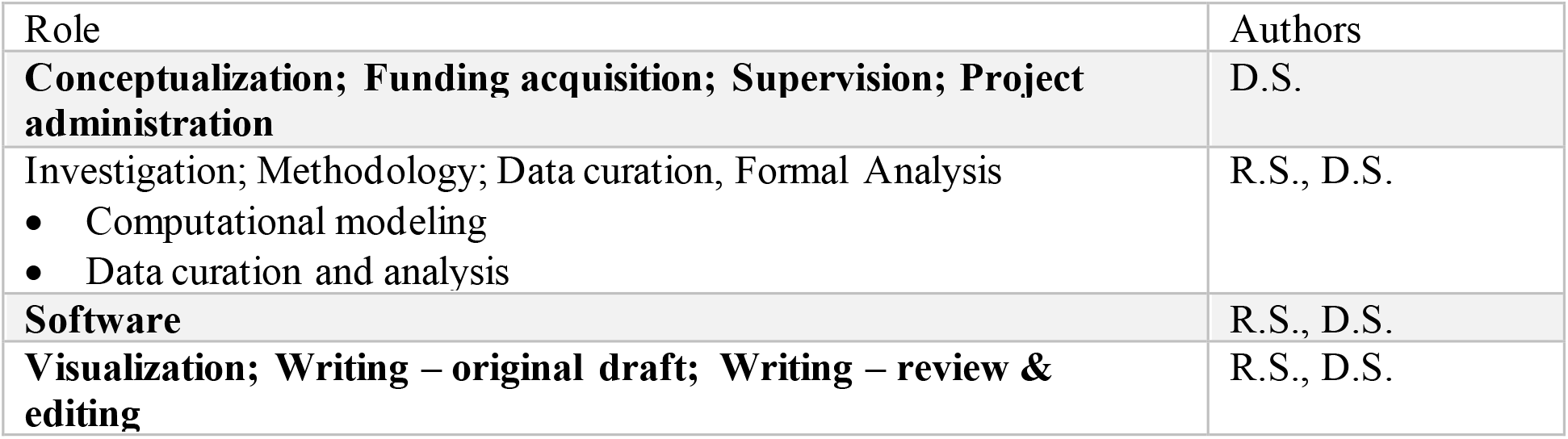

## Acknowledgements

This work was supported by the National Institutes for Health (NIH) grants R00-CA151673, R01-GM138385, UG3 TR003355, R01-AI155696 (to DS), UCOP-RGPO (R00RG2628 & R00RG2642 to DS), The Sanford Stem Cell Clinical Center at UCSD (to DS), Padres Pedal the Cause / Rady Children’s Hospital Translational PEDIATRIC Cancer Research Award (Padres Pedal the Cause/RADY #PTC2017) to DS, 2017, Padres Pedal the Cause /C3 Collaborative Translational Cancer Research Award (San Diego NCI Cancer Centers Council (C3) #PTC2017) to DS.

## Data and materials availability

All data is available in public repository and the relevant accession number are provided in the text and the supplementary materials.

---

**Figure S1.**
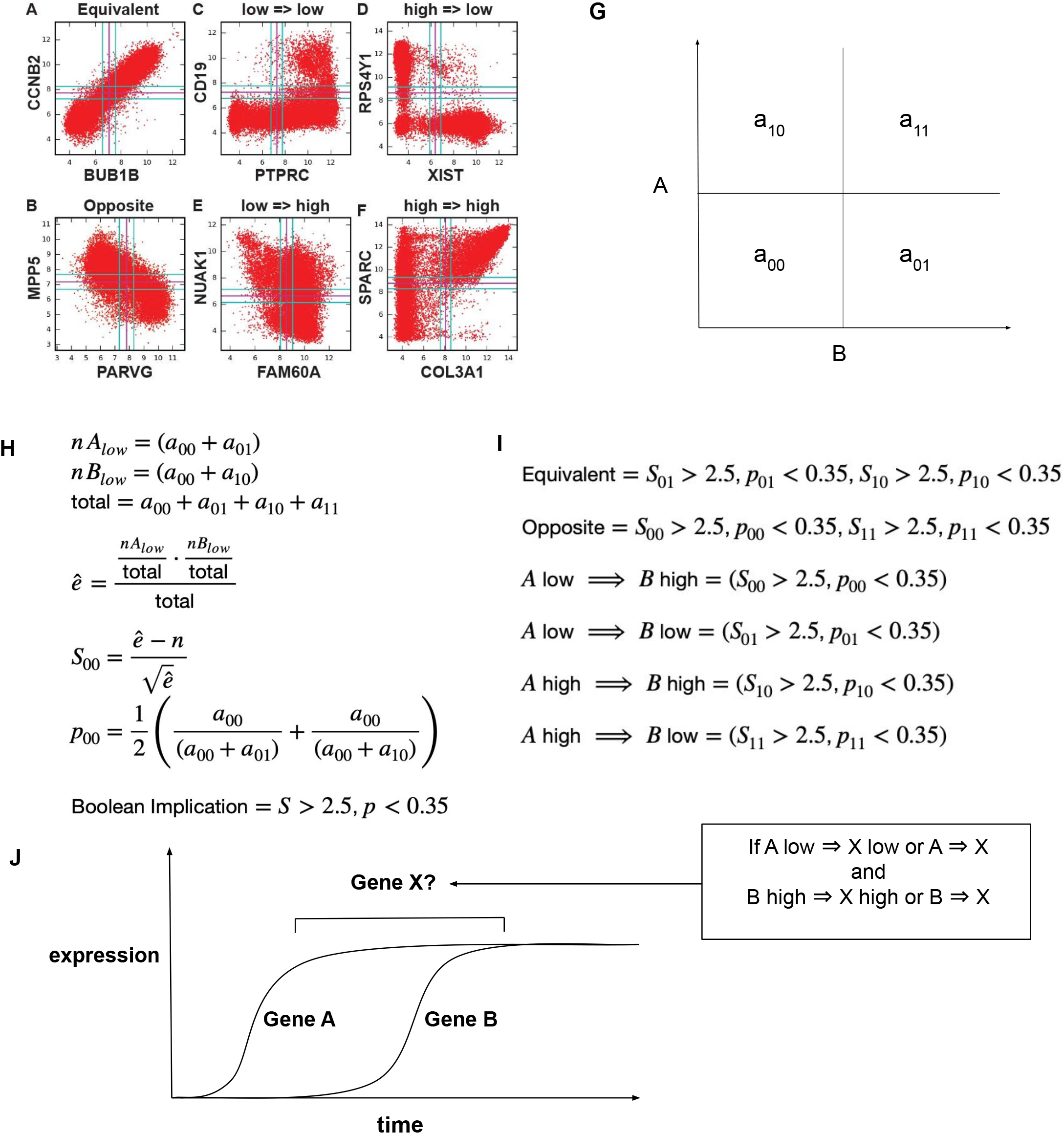
Discovery of Boolean Implication Relationships. Method for discovering and applying Boolean implication relationships in single cell RNA sequencing data. **(A-F):** Six types of Boolean implication relationships are visible on scatter plots. Two are symmetric with two sparse quadrants (A-B) and four are asymmetric with one sparse quadrant (C-F). **(G):** This plot is divided into four quadrants based on thresholds identified by the StepMiner algorithm. **(H-I):** The BooleanNet algorithm identifies the sparse quadrants using a statistic *S* and a likelihood error rate *p* and applying thresholds of 2.5 and 0.35, respectively. **(J):** Analysis of Boolean implication relationships was used to find genes involved in cell fate determination using bait genes (A and B).

**Figure S2.**
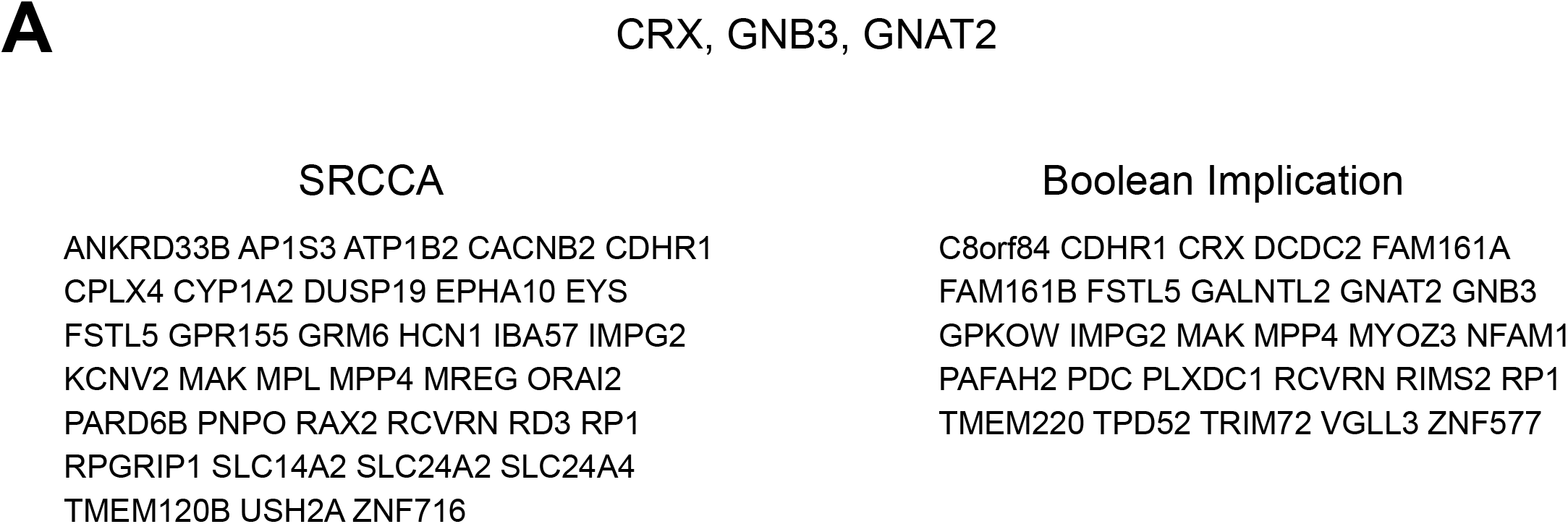
Additional Gene Lists. **(A):** Additional gene lists from SRCCA and Boolean analysis using common bait genes for cones CRX, GNAT2 and GNB3.

